# Pooled CAR-T screening in nonhuman primates identifies designs with enhanced proliferation, trafficking, and persistence

**DOI:** 10.1101/2025.03.05.640197

**Authors:** Lucy H. Maynard, Eric J. Cavanaugh, Haiying Zhu, Carly E. Starke, Sarah M. Doherty, Teresa Einhaus, Ailyn Perez, Laurence Stensland, Michelle Hoffman, Veronica Nelson, Sarah Herrin, Chad Littlewood, Kaycee Camou, Erica Wilson, Christopher Wessel, Keith R. Jerome, Hans-Peter Kiem, Christopher W. Peterson

## Abstract

Chimeric antigen receptor T (CAR-T) cell therapy has revolutionized treatment for B-cell malignancies, yet over 60% of patients relapse within one year, often due to insufficient CAR-T persistence. While mouse and primary cell models have been instrumental in advancing CAR-T therapy, they frequently fail to predict clinical outcomes, underscoring the need for more translationally relevant models. To address this limitation, we conducted the first systematic evaluation of CAR structure-function relationships in an immunocompetent nonhuman primate (NHP) model. We engineered an array of 20 CD20-targeted CARs with distinct combinations of hinge, transmembrane, and costimulatory domains. Following *ex vivo* characterization, we administered pooled autologous CAR-T arrays to three NHPs and tracked CAR abundance longitudinally using a novel digital droplet PCR assay. *Ex vivo*, CAR-T cells incorporating the MyD88-CD40 costimulatory domain exhibited markedly distinct functional profiles, including increased activation, unique cytokine secretion, tonic signaling, and resistance to exhaustion. *In vivo*, MyD88-CD40 CARs expanded dramatically, comprising up to 100% of peripheral T cells and significantly outperforming canonical CD28- and 4-1BB-based CARs. This expansion was associated with robust B-cell depletion across all animals. MyD88-CD40 CARs, particularly those with a CD28 hinge and transmembrane domain, demonstrated superior trafficking to secondary lymphoid tissues and persistence through study endpoint, unlike other CARs which waned by day 28. Our findings highlight the value of NHP models for screening CAR designs and identify MyD88-CD40 CARs as candidates with unmatched potency. The unique functional attributes conferred by this domain may provide key insights into features that drive enhanced CAR-T cell activity.

**Key points:**

- We developed the first pooled CAR-T screening platform in an immunocompetent nonhuman primate model to directly compare CAR designs.
- We identified MyD88-CD40 costimulatory domain as vastly superior to conventional domains in proliferation, trafficking and persistence.

## INTRODUCTION

Chimeric antigen receptor T (CAR-T) cell therapy has transformed the treatment landscape for B-cell malignancies, leading to the approval of eight therapies targeting CD19 and B-cell maturation antigen (BCMA). However, tumor recurrence and patient relapse remain significant challenges. Relapse occurs in 50-70% of patients with diffuse large B-cell lymphoma (DLBCL) and B-acute lymphoblastic leukemia (B-ALL) treated with CD19-targeted CAR-T cells,^1–6^ and in over 80% of multiple myeloma (MM) patients receiving BCMA-targeted CAR-T cells.^7,8^ A meta-analysis spanning eight clinical studies across nine B-cell malignancies further underscores this issue, revealing that over 60% of these patients relapse within just 12 months after CAR-T infusion, emphasizing the rapid progression of disease recurrence.^9^

The high rate of relapse highlights a critical need to improve the long-term efficacy of CAR-T cell therapy. While various factors contribute to relapse, mounting evidence suggests that limited *in vivo* persistence of CAR-T cells is a primary challenge.^10^ Relapse of antigen-positive tumor cells is frequently associated with limited CAR-T durability,^11,12^ detected in 33-78% of CD19-targeted^4,11–14^ and 67-96% of BCMA-targeted CAR-T cell^7,15,16^ relapse cases. Recent studies have further identified CAR-T cell persistence as a crucial determinant of long-term disease-free survival,^17,18^ highlighting the need for strategies to enhance CAR-T durability.

The structure of a CAR plays a critical role in shaping CAR-T cell function, phenotype, and persistence.^19^ CAR molecules comprise several components, including antigen recognition, hinge, transmembrane, costimulatory, and CD3ζ activation domains. The costimulatory domain is particularly influential in modulating CAR-T cell kinetics and durability.^20,21^ Approved CAR-T therapies incorporate either 4-1BB or CD28 costimulatory domains, which exhibit distinct profiles: 4-1BB promotes greater persistence with slower expansion, while CD28 drives rapid expansion but earlier T-cell exhaustion.^22,23^ Recent studies have identified additional domains, such as Ox40,^24–26^ BAFF-R,^27^ and MyD88-CD40,^28–30^ which have demonstrated the potential to further augment CAR-T cell activity. Hinge and transmembrane domains also influence CAR-T cell activity, particularly by modulating sensitivity to antigen levels.^31–35^ Despite these advances, no universally superior CAR architecture has been identified, suggesting a requirement for disease-specific designs. As such, direct head-to-head comparisons of CAR structures could provide critical insights into designs that improve CAR-T cell activity for specific indications.

Developing and validating novel CAR designs through clinical trials is an exceedingly expensive, time-consuming, and resource-intensive process, with a substantial risk of failure. Moreover, commonly used mouse models frequently fail to recapitulate complex human immunology and CAR-T-mediated toxicities, such as cytokine release syndrome (CRS), limiting their utility in predicting clinical outcomes.^36,37^ To address these challenges, the nonhuman primate (NHP) model has emerged as a valuable tool for studying CAR-T therapy. Unlike immunocompromised mouse models, NHPs possess a functional immune system and can recapitulate drug-associated toxicities observed in humans. Furthermore, the longer lifespan of NHPs allows for extended safety and efficacy studies, and their larger size enables longitudinal biopsies and testing of clinical-scale CAR-T manufacturing, which are not feasible in mice.^38^ Our research group and others have previously developed and validated a NHP model for B-cell targeted CAR-T cell therapy.^39–42^ This model has demonstrated its utility in recapitulating key clinical findings, including potent B-cell depletion in blood and tissues, transient ablation of B-cell follicles, and management of CAR-T cell-mediated toxicities. Here, we hypothesized that the NHP model could be leveraged in a pooled screening format to simultaneously evaluate multiple CAR structures and identify designs with superior *in vivo* activity. Our findings underscore the importance of CAR design in shaping CAR-T cell proliferation, persistence, and trafficking, and highlight the value of immunocompetent NHP models for advancing next-generation CAR-T therapies.

## METHODS

### CAR-T cell design and manufacturing

The CAR array was designed to span all combinations of hinge (CD8, CD28), transmembrane (CD8, CD28), and costimulatory (4-1BB, CD28, BAFF-R, MyD88-CD40, Ox40) domains, with a common anti-CD20 scFv and CD3ζ activation domain (further details in Supplemental Methods). Lentiviral vectors were generated using a third-generation packaging system pseudotyped with Vesicular Stomatitis Virus G-protein (Fred Hutch Preclinical Vector Core), and titered via qPCR in HT1080 cells.^43^ CAR-T cells were manufactured *ex vivo* as described.^44^ Briefly, CD4^+^ and CD8^+^ T cells were isolated from peripheral blood mononuclear cells (PBMCs) using bead-based selection (STEMCell Technologies) and cultured in X-VIVO-15 (Thermo Fisher Scientific) supplemented with 10% FBS, 1% penicillin/streptomycin, 1% L-glutamine, 55 μM 2-mercaptoethanol, and 5 ng/mL recombinant IL-7 and IL-15 (Peprotech). T cells were activated with artificial antigen-presenting cells (aAPCs; kind gift from Dr. James Riley^45^) at a 2:1 ratio. Three days later, T cells were transduced with lentiviral vectors at a multiplicity of infection from 1–10 with 5 μL of LentiBlast Transduction Reagent (OZ Bioscience), followed by 7-day expansion in G-REX24 flasks (Wilson Wolf).

### CellTrace Violet Proliferation Assay

CAR-T cell proliferation was assessed immediately after manufacturing using CellTrace Violet dye (CTV, Thermo Fisher Scientific). CAR-T cells were CTV-labeled according to manufacturer instructions, and 5×10⁵ labeled CAR-T cells were co-cultured with 1×10⁵ irradiated CD20- expressing K562 cells (K562-CD20, kindly provided by Dr. Brian Till^46^) or wild-type K562 cells (K562-wt; ATCC) (effector-to-target ratio 5:1) for three days at 37°C. Proliferation was quantified by flow cytometry based on CTV dilution in CAR^+^ and CAR^-^ subsets.

### Real-time cell analysis of cytotoxicity

CAR-T cell cytotoxicity was assessed using the xCELLigence (Agilent Technologies) real-time cell analysis (RTCA) platform. CAR-T cells were co-cultured with CD20-expressing LLCMK2 (LLCMK2-CD20)^40^ or wild-type LLCMK2 (LLCMK2-wt; ATCC) target cells at varying effector-to-target ratios, and target cell death was quantified as percent cytolysis (%). Cytotoxicity was measured immediately after manufacturing and following six rounds of stimulation with irradiated K562-CD20 cells.

### Flow cytometry and chronic antigen stimulation assay

All antibodies and clones for flow cytometry are listed in **Supplemental Table 2**. All flow panels included a live/dead stain to exclude dead cells. The long-term function of CAR-T cell variants was evaluated through serial stimulation with irradiated K562-CD20 cells. To standardize initial conditions, CAR-T variants were analyzed for CAR expression via flow cytometry and diluted to 20% CAR^+^ cells by adding untransduced autologous T cells. 5×10^5^ CAR-T cells were co-cultured with 1×10^5^ irradiated K562-CD20 or K562-wt cells (effector-to-target ratio of 5:1). Every 3-4 days, CAR-T cells were harvested, with half being restimulated and half stained with immunophenotyping panels and analyzed by flow cytometry. The same experiment was performed with equal numbers of pooled CAR^+^ cells from each CAR-T variant, with half being restimulated and half snap-frozen for analysis via ddPCR.

### Cytokine measurements

NHP plasma samples and CAR-T cell supernatants collected 24 hours after co-culture with K562-CD20 or K562-wt cells were assayed using the LEGENDplex™ 10-plex NHP Th Cytokine Panel (BioLegend). Cerebrospinal fluid (CSF) cytokines were measured using the ProcartaPlex™ 14-plex NHP Th Panel (Invitrogen), following the manufacturer’s instructions.

### *In vivo* NHP CAR-T cell studies

This study was approved by the Institutional Animal Care and Use Committees of the Fred Hutchinson Cancer Center/University of Washington (protocol no. 3235-04). Pigtail macaques were housed and cared for under conditions that meet NIH standards as stated in the *Guide for the Care and Use of Laboratory Animals* (National Research Council, National Academy Press, Washington, DC, 1996), ILAR recommendations, and AAALAC accreditation standards, as described previously.^44^ Animals received cyclophosphamide (40 mg/kg) on days -7 and -6 prior to CAR-T infusion. To ensure equal pooling of CAR variants in the infusion product, CAR-T variants were analyzed for CAR expression by flow cytometry, and equal numbers of CAR^+^ variants were pooled to achieve a final dose of 9×10^6^ CAR^+^ cells/kg. Irradiated K562-CD20 cells were infused at 2.5×10^7^ cells/kg on days 21 and 35 following CAR-T infusion for antigen restimulation, as previously described.^44^ Blood was collected longitudinally until necropsy. Sampling in other compartments including spleen, lymph nodes, CSF, bone marrow and bronchoalveolar lavage (BAL) was performed at -2, 2 and 4 weeks following CAR-T infusion. Tissues were processed and dissociated for downstream analyses using established protocols.^40^ Details on CRS management and immunohistochemistry are presented in the Supplemental Methods.

### Droplet digital PCR assay

Droplet digital PCR (ddPCR) was used to quantify CAR variant abundance. Unique primer/probe sets were designed and validated for specific detection. Genomic DNA (gDNA) was extracted (QIAGEN; AllPrep DNA), diluted to ∼100 ng/µL, and divided across six ddPCR assays, with 1-10 µL of gDNA per reaction. Each multiplex assay targeted a subset of CAR variants (2- to 6-plex). To normalize these data, 1:10 diluted gDNA was used to detect a housekeeping gene (macaque RNAse-P p30 subunit, MRPP30)^47^ and T cell receptor D (TRD), a gene that is excised during T cell receptor gene rearrangement and is therefore only present in non-T cells.^48^ The ddPCR reactions were performed using the Droplet Digital PCR System (Bio-Rad) with ddPCR Multiplex Supermix reagents. Negative controls (untransduced T cells, H_2_0) were routinely included to ensure the specificity of amplification. Samples with >3 ddPCR-positive droplets were considered positive. Further information about ddPCR normalization and assay sensitivity measurements are presented in the Supplemental Methods.

## RESULTS

### Design and characterization of a combinatorial CD20-targeted CAR array

To systematically assess how CAR structure impacts CAR-T cell activity, we designed an array of CD20-directed CARs, termed V1-V20, spanning all combinations of established hinge (CD8, CD28), transmembrane (CD8, CD28), and costimulatory domains (4-1BB, CD28, BAFF-R, MyD88-CD40, Ox40) (**Figure 1A and Table 1**). To enable direct comparison, we retained the same validated CD20 scFv and CD3ζ activation domains across all designs,^39,40^ and included a co-expressed epidermal growth factor receptor (EGFRt) to detect CAR-transduced cells.^49^ We manufactured the CAR-T cell array using established methods,^39,40,44^ including separate isolation and activation of CD4^+^ and CD8^+^ cells from PBMCs, transduction with lentiviral vectors, and expansion in flasks designed to maximize gas and nutrient convection (**Figure 1B**).

**Figure 1.**
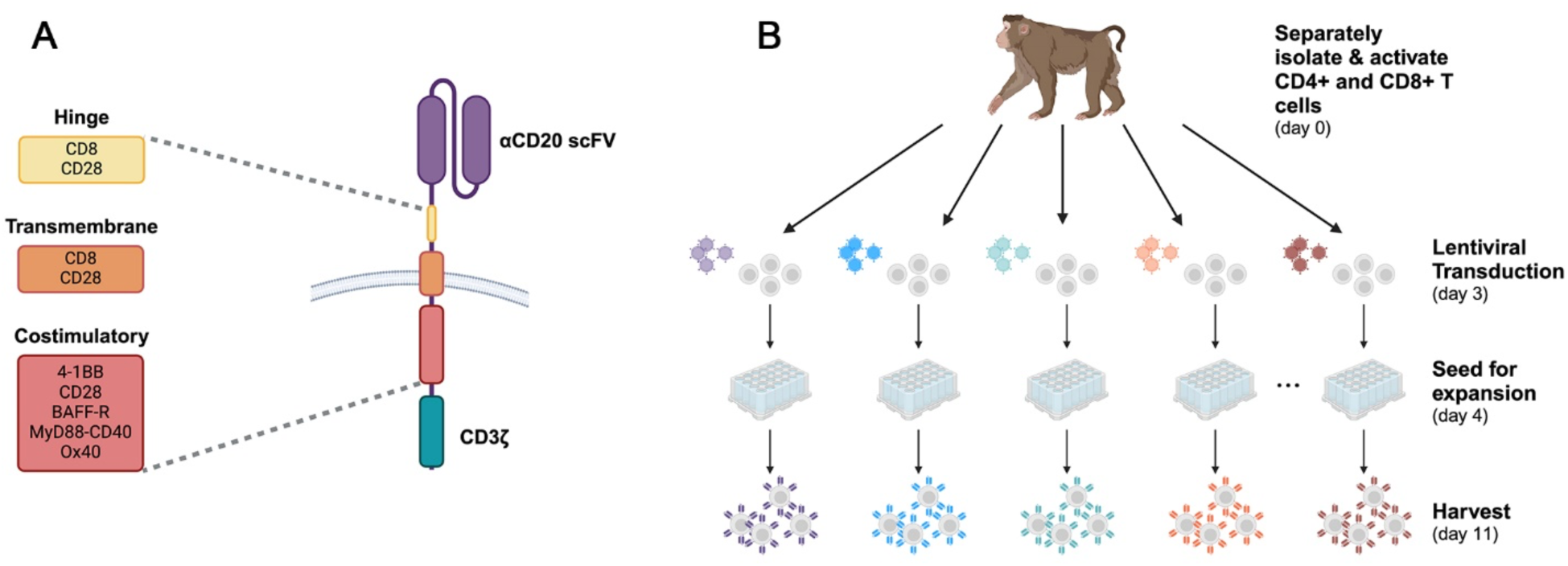
Design and generation of a combinatorial array of CD20-targeted CAR-T cells. (A) An array of chimeric antigen receptor (CAR) constructs was engineered to systematically explore the impact of 20 distinct CAR architectures in the same animal. The array spans all combinations of hinge (CD8 or CD28), transmembrane (CD8 or CD28) and costimulatory (4-1BB, CD28, BAFF-R, MyD88-CD40, Ox40) domains. (B) CAR-T cells were manufactured *ex vivo*: autologous CD4^+^ and CD8^+^ cells from pigtail macaques (*M. nemestrina*) were isolated, activated, transduced with CAR-encoding lentiviral vectors, and expanded before harvesting for infusion and functional assays.

**Table 1.**
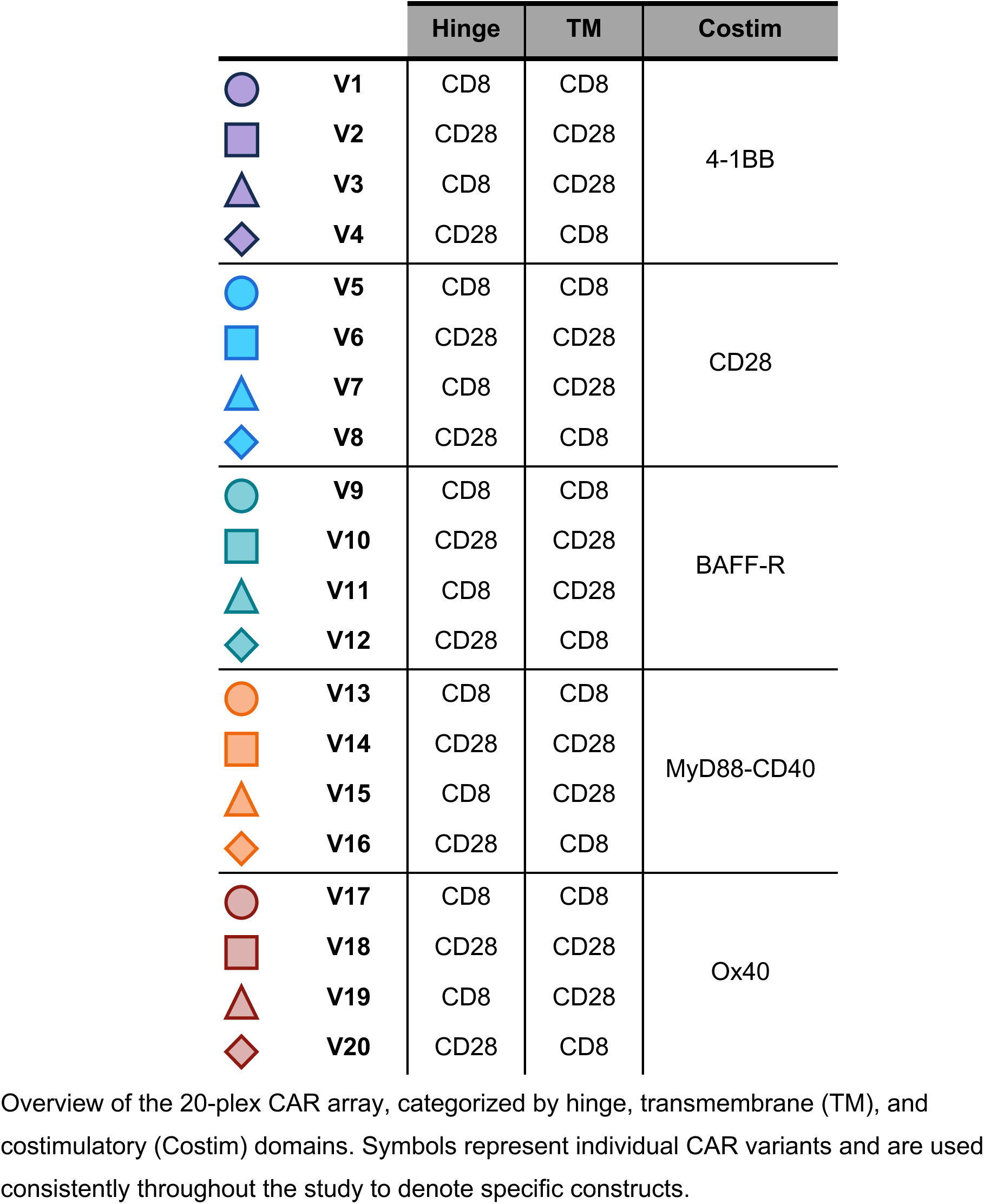
Domain structures for the combinatorial CAR array.

|  |  | Hinge | TM | Costim |
| --- | --- | --- | --- | --- |
|  | V1 | CD8 | CD8 | 4-1BB |
|  | V2 | CD28 | CD28 |  |
|  | V3 | CD8 | CD28 |  |
|  | V4 | CD28 | CD8 |  |
|  | V5 | CD8 | CD8 | CD28 |
|  | V6 | CD28 | CD28 |  |
|  | V7 | CD8 | CD28 |  |
|  | V8 | CD28 | CD8 |  |
|  | V9 | CD8 | CD8 | BAFF-R |
|  | V10 | CD28 | CD28 |  |
|  | V11 | CD8 | CD28 |  |
|  | V12 | CD28 | CD8 |  |
|  | V13 | CD8 | CD8 | MyD88-CD40 |
|  | V14 | CD28 | CD28 |  |
|  | V15 | CD8 | CD28 |  |
|  | V16 | CD28 | CD8 |  |
|  | V17 | CD8 | CD8 | Ox40 |
|  | V18 | CD28 | CD28 |  |
|  | V19 | CD8 | CD28 |  |
|  | V20 | CD28 | CD8 |  |

Overview of the 20-plex CAR array, categorized by hinge, transmembrane (TM), and costimulatory (Costim) domains. Symbols represent individual CAR variants and are used consistently throughout the study to denote specific constructs.

All CAR variants were successfully manufactured *ex vivo*, with MyD88-CD40 CAR-T cells exhibiting enhanced expansion during manufacturing (**Figure 2A**). Each variant demonstrated robust CAR expression (**Figure 2B**), balanced CD4:CD8 ratios (**Figure 2C**), and a predominant central memory phenotype (CD28^+^ CD95^+^; **Figure 2D**).^39^ Among these, BAFF-R and MyD88-CD40 CAR-T cells displayed higher CD4:CD8 ratios and a greater proportion of central memory T-cells compared to 4-1BB or CD28 CAR-T cells. Notably, MyD88-CD40 CAR-T cells showed significantly increased expression of CD25 (**Figure 2E**, *P* < 0.0005) and HLA-DR (**Figure 2F**, *P* < 0.00005) activation markers,^50^ suggesting this domain confers an hyperactivation phenotype. These data validate our arrayed anti-CD20 CAR-T cell manufacturing and highlight CAR-dependent phenotypes that emerge during *ex vivo* manufacturing.

**Figure 2.**
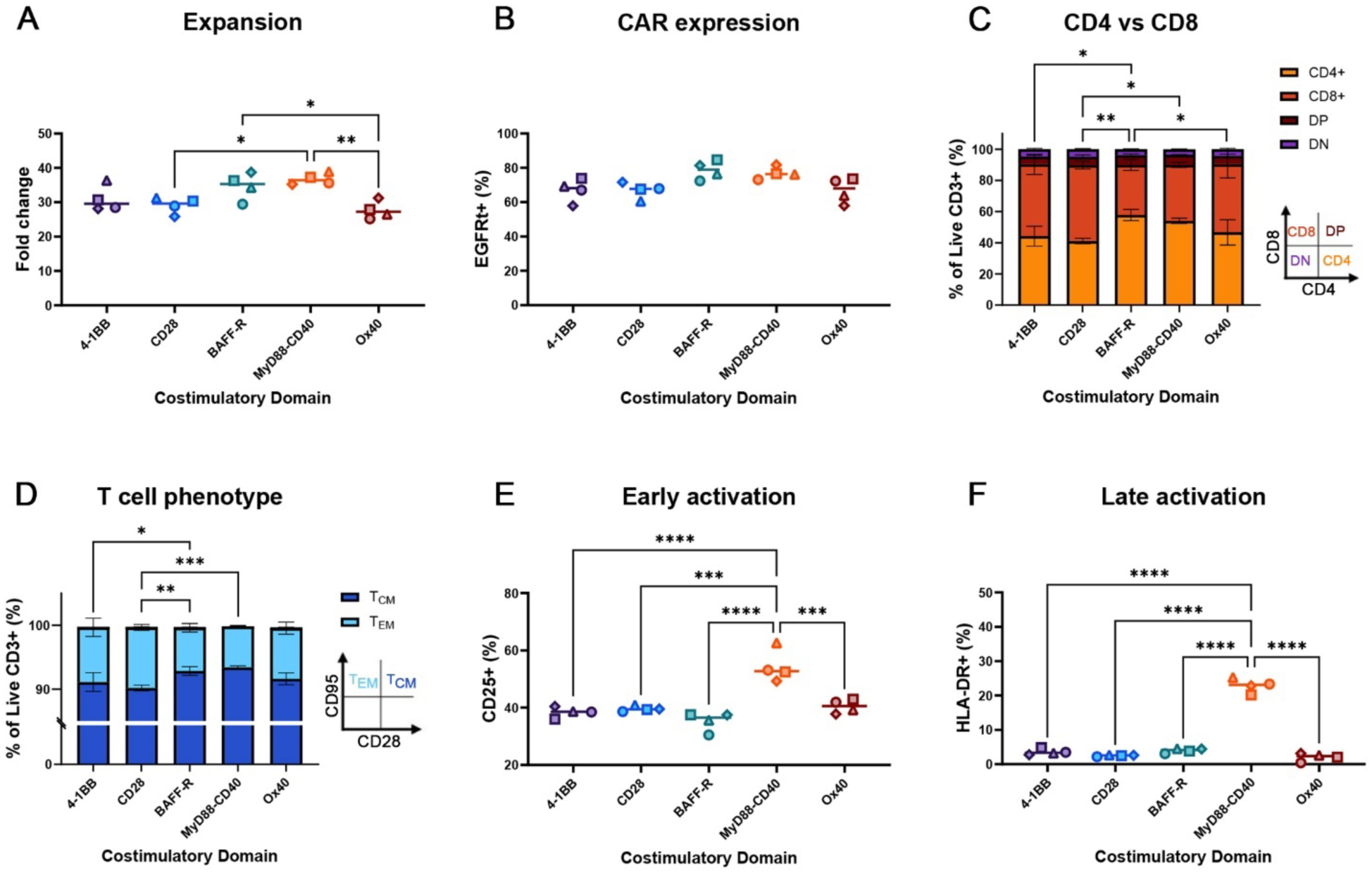
Phenotypic characterization of individual CAR-T variants. (A) Following arrayed *ex vivo* manufacturing, CAR-T cell variants V1-V20 were harvested and counted. Fold change represents change in total cell counts over 7 days of expansion in G-REX flasks. (B-F) At harvest, each CAR-T variant was assessed via flow cytometry for (B) CAR expression via truncated EGFR (EGFRt); (C) CD4/CD8 T cell composition including CD4^+^ single positive (CD4^+^ CD8^-^), CD8^+^ single positive (CD4^-^ CD8^+^), double positive (DP, CD4^+^ CD8^+^), and double negative (DN, CD4^-^ CD8^-^); (D) T cell subsets, T_CM_ (Central memory, CD28^+^ CD95^+^), T_EM_ (Effector memory, CD28^-^ CD95^+^); (E) early T cell activation marker CD25 in EGFRt^+^ subsets; and (F) late T cell activation marker HLA-DR in EGFRt^+^ subsets. Each datapoint represents the average from three biological replicates (*n* = 3 pigtail macaques), except for V17 which represents the average from two biological replicates (*n* = 2 pigtail macaques). Statistical significance was determined using one-way ANOVA with Tukey’s correction for multiple comparisons. \**P* < 0.05, \*\**P* < 0.005, \*\*\**P* < 0.0005, \*\*\*\**P* < 0.00005. In panels A, B, E, and F, shapes and color codes correspond to Table 1.

### CAR-T variants display distinct functional profiles *ex vivo*

We next extended our *ex vivo* experiments to explore the impact of CAR architecture on CAR-T cell function, including proliferative, cytokine, and cytolytic responses. To measure CAR-specific proliferation, we quantified the dilution of CellTrace Violet (CTV) dye (**Figure 3A**). We observed robust division of CAR^+^ cells when stimulated with K562-CD20 targets, with minimal bystander proliferation in the CAR^-^ fraction (**Figure 3B**, *P* < 0.00005). Notably, MyD88-CD40 CAR-T cells uniquely exhibited significant tonic signaling, as indicated by antigen-independent proliferation of CAR^+^ cells (**Figure 3C**, *P* < 0.00005). These cells displayed phenotypic features of tonic signaling,^51^ including elevated CD25 and HLA-DR, as well as reduced expression of exhaustion markers PD-1 and TIGIT (**Figure 3D**). Following K562-CD20 stimulation, all CAR variants exhibited antigen-specific cytokine secretion (**Supplemental Figure 1A-G**), with distinct patterns across costimulatory domain groups. Most notably, CD20-stimulated MyD88-CD40 CAR-T cells produced minimal IL-6 (**Figure 3E**), significantly reduced IL-10 (**Figure 3F**, *P* < 0.005), and increased IL-13 (**Figure 3G**, *P* < 0.005), a cytokine less characterized in CAR-T cell biology but previously linked to MyD88-CD40 signaling.^29^ Assays measuring the specific lysis of target cells over time (**Figure 3H**), confirmed antigen-specific cytolysis of LLCMK2-CD20 target cells across all CAR variants, with minimal non-specific killing of LLCMK2-wt cells (**Figure 3I**, *P* < 0.00005). Interestingly, MyD88-CD40 CAR-T cells also displayed modest-to-significantly enhanced antigen-dependent cytotoxicity (**Supplemental Figure 1H**). Finally, we applied principal component analysis (PCA) to integrate data from functional assays (**Figure 3B-I**). PCA further highlighted the distinctiveness of MyD88-CD40 CAR-T cells, which clustered separately from other variants (**Figure 3J**). Collectively, these findings underscore the influence of costimulatory domains on CAR-T function, with MyD88-CD40 CARs distinguished by tonic signaling, unique cytokine profiles, and potent cytolytic activity.

**Figure 3.**
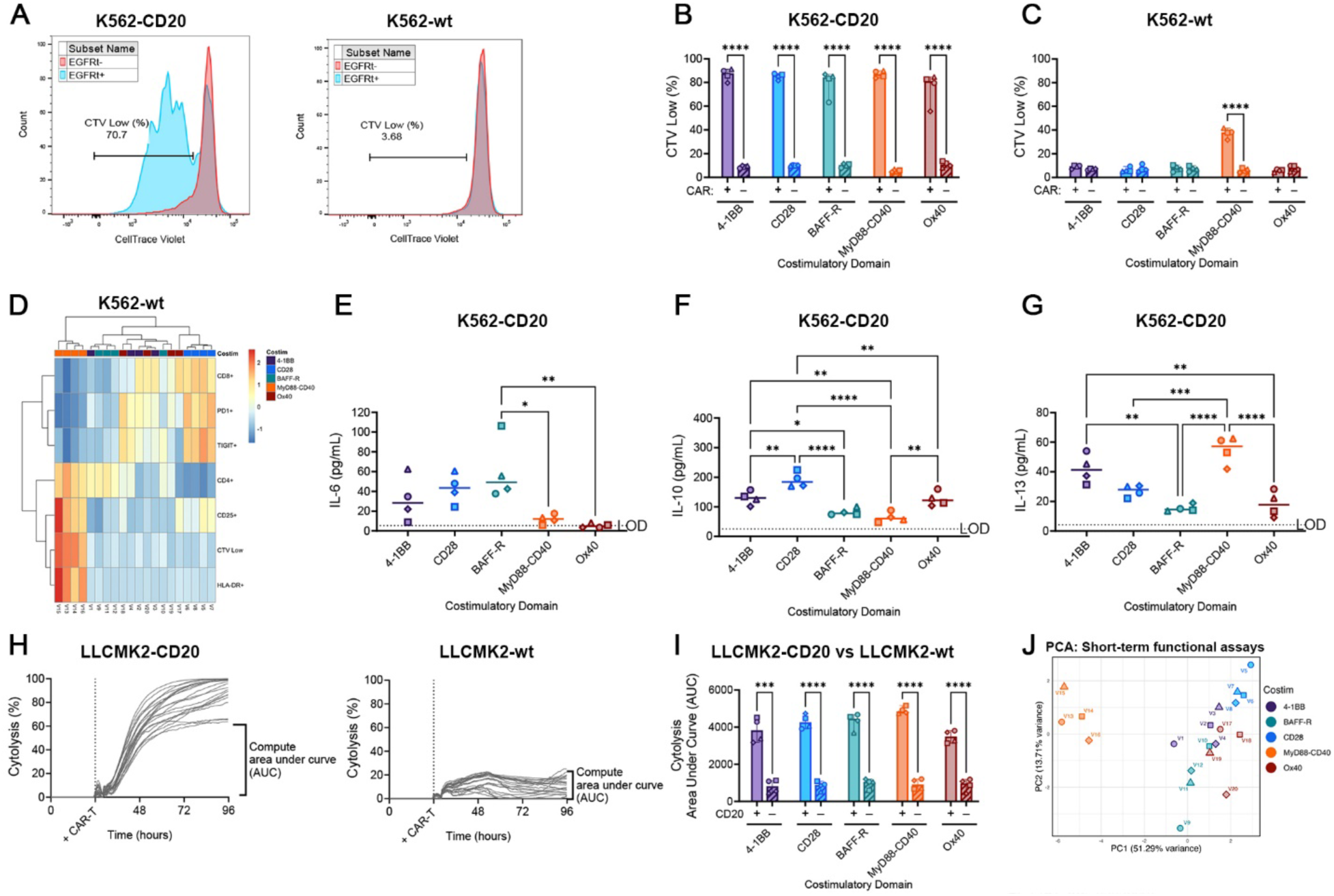
CAR-T cell variants exhibit distinct functional profiles under antigen-positive and antigen-negative stimulation. (A) Proliferation was assessed by quantifying dilution of CellTrace Violet (CTV Low) in CAR^+^ (EGFRt^+^) and CAR^-^ (EGFRt^-^) subsets. (B-C) quantification of CTV Low cells following co-culture with irradiated (B) antigen-positive K562-CD20 and (C) antigen-negative wild-type K562 cells. CAR^+^ and CAR^-^ subsets are indicated on the x-axis. (D) Z-normalized flow cytometry data for CAR^+^ subsets stimulated with K562-wt cells. (E-G) Cytokine secretion for (E) IL-6, (F) IL-10, and (G) IL-13 was measured 24 hours after co-culture with K562-CD20 cells. (H) Cytotoxicity was tracked using the xCELLigence RTCA assay, measuring target cell lysis over time. (I) Area under the curve (AUC) analysis of cytolysis curves for antigen-positive (LLCMK2-CD20) and antigen-negative (LLCMK2-wt) target cells from (H). Antigen status (+ or -) is denoted on the x-axis. (J) Data from (B-I) was compiled, Z-normalized and analyzed by principal component analysis (PCA). For (A-I), the antigen status of the target cells (e.g., K562-CD20, LLCMK2-wt) is indicated above each graph. Data represent the average of three biological replicates (*n* = 3 pigtail macaques) for (B-G), except for V17 (*n* = 2). Two replicates were used for (H) and (I). Statistical significance was determined using multiple unpaired T-tests with Holm-Šídák correction (B, C, I) and one-way ANOVA with Tukey’s correction (E-G). \**P* < 0.05, \*\**P* < 0.005, \*\*\**P* < 0.0005, \*\*\*\**P* < 0.00005.

### Chronic antigen stimulation highlights phenotypic and functional differences among CAR-T cell variants

To investigate the influence of CAR design on *in vivo* persistence,^22,23^ we next compared the long-term function of CAR-T variants in an *ex vivo* chronic antigen stimulation assay. CAR-T cells were repeatedly exposed to K562-CD20 cells to evaluate CAR-T durability amid exhaustive conditions (**Figure 4A**).^52–55^ All variants demonstrated robust expansion, marked by increased EGFRt^+^ expression across successive stimulations (**Figure 4B**). Consistent with previous reports,^28,29,56^ only MyD88-CD40 CARs retained lower PD-1 (**Figure 4C**, *P* < 0.0005) and TIGIT (**Figure 4D**, *P* < 0.005) after chronic stimulation, suggesting this domain confers superior resistance to exhaustion. MyD88-CD40 CAR-T cells also produced less IL-10 (**Figure 4E**, *P* < 0.005) and substantially more IL-13 (**Figure 4F**, *P* < 0.00005). Interestingly, 4-1BB CAR-T cells failed to retain antigen-specific cytotoxicity after chronic stimulation (**Figure 4G**). PCA of chronic stimulation data further highlighted the distinct clustering of MyD88-CD40 and 4-1BB CARs (**Figure 4H**), underscoring the role of costimulatory domains in shaping CAR-T exhaustion, cytokine secretion, and cytotoxicity during prolonged antigen exposure.

**Figure 4.**
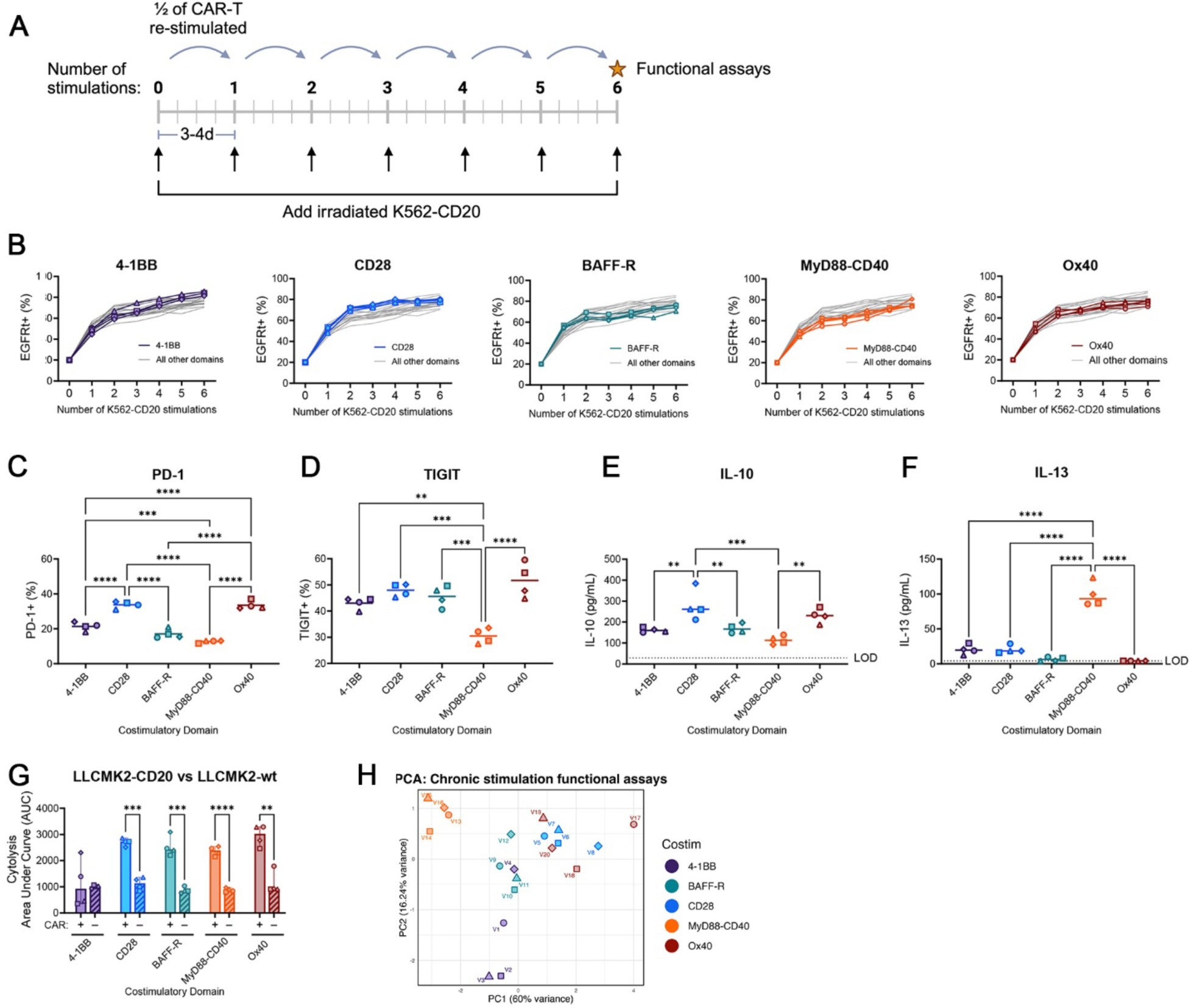
Chronic antigen stimulation *ex vivo* reveals functional divergence between CAR-T cell variants. (A) *Ex vivo*-manufactured CAR-T cells were quantified and normalized to 20% CAR^+^ by addition of untransduced T cells, then repeatedly stimulated with irradiated K562-CD20 every 3–4 days. Over six rounds of stimulation, cells were harvested for immunophenotyping and half were restimulated. After the final round of stimulation, cells were assessed for cytotoxicity and cytokine secretion. (B) CAR expression levels; each panel highlights one of 5 costimulatory domain groups. (C-D) Expression of exhaustion markers (C) PD-1 and (D) TIGIT by flow cytometry. (E-F) Secretion of (E) IL-10 and (F) IL-13 following chronic stimulation. (G) Cytolysis AUC for LLCMK2-CD20 and LLCMK2-wt target cells after chronic stimulation. (H) Data from (C-G) was compiled, Z-normalized and analyzed via PCA. Each datapoint in (B-F) and LLCMK2-CD20 cytolysis in (G) represents the average of three biological replicates (*n* = 3 pigtail macaques), except for V17 (*n* = 2). LLCMK2-wt cytolysis in (G) represents two biological replicates. Statistical significance was determined using one-way ANOVA with Tukey’s correction for (C-F) and multiple unpaired T-tests with Holm-Šídák correction for (G). \**P* < 0.05, \*\**P* < 0.005, \*\*\**P* < 0.0005, \*\*\*\**P* < 0.00005.

### Pooled chronic stimulation *ex vivo* reveals donor-specific proliferation of CAR-T cell variants

Prior to *in vivo* experiments where CAR variants were pooled and infused into single recipient NHPs, we developed a competitive *ex vivo* stimulation assay in which pooled CAR-T variants were serially stimulated with K562-CD20 as in **Figure 4A**. Since flow cytometry cannot distinguish unique CAR variants, we turned to CAR quantitation at the DNA level using ddPCR. Our variant-specific ddPCR assay was designed to enable precise quantification of individual CAR variants within pooled samples from *ex vivo* and *in vivo* experiments. Primer-probe sets were designed for individual CAR variants and validated to ensure variant-specific amplification (**Supplemental Figure 2A**), and assay sensitivity to single-copy thresholds (**Supplemental Figure 2B**). Donor-specific expansion patterns emerged, with 4-1BB or CD28 CARs preferentially expanding (**Supplemental Figure 3**). Notably, V7 (CD8 hinge, CD28 transmembrane, CD28 costimulatory domain), showed significant enrichment in two donors, whereas BAFF-R or MyD88-CD40 CARs exhibited no expansion across all donors. These data suggest that CD28 and 4-1BB costimulatory domains promote efficient proliferation in competitive *ex vivo* assays. We next performed a corresponding competitive *in vivo* study using the same biological donors to determine whether these *ex vivo* findings were recapitulated *in vivo*.

### Pooled CAR-T cells expand *in vivo*, induce B-cell aplasia, and exhibit distinct inflammatory profiles

Following extensive *ex vivo* characterization, we next evaluated our CAR array in a cohort of three pigtail macaques. We pooled CAR-T variants in equal proportions (**Supplemental Figure 4**) and infused into autologous NHPs preconditioned with a low-dose cyclophosphamide regimen.^40^ Each animal received a dose of 9×10⁶ CAR^+^ cells/kg (**Figure 5A and Supplemental Table 1**). To align with our *ex vivo* data and methods in prior studies from our group and others,^39,44^ we provided supplemental antigen in the form of irradiated K562-CD20 cells that were administered at 3 and 5 weeks post-CAR infusion. CAR-T proliferation, trafficking, and persistence were evaluated in longitudinal blood and tissues for up to 102 days post-CAR infusion (**Figure 5B**).

**Figure 5.**
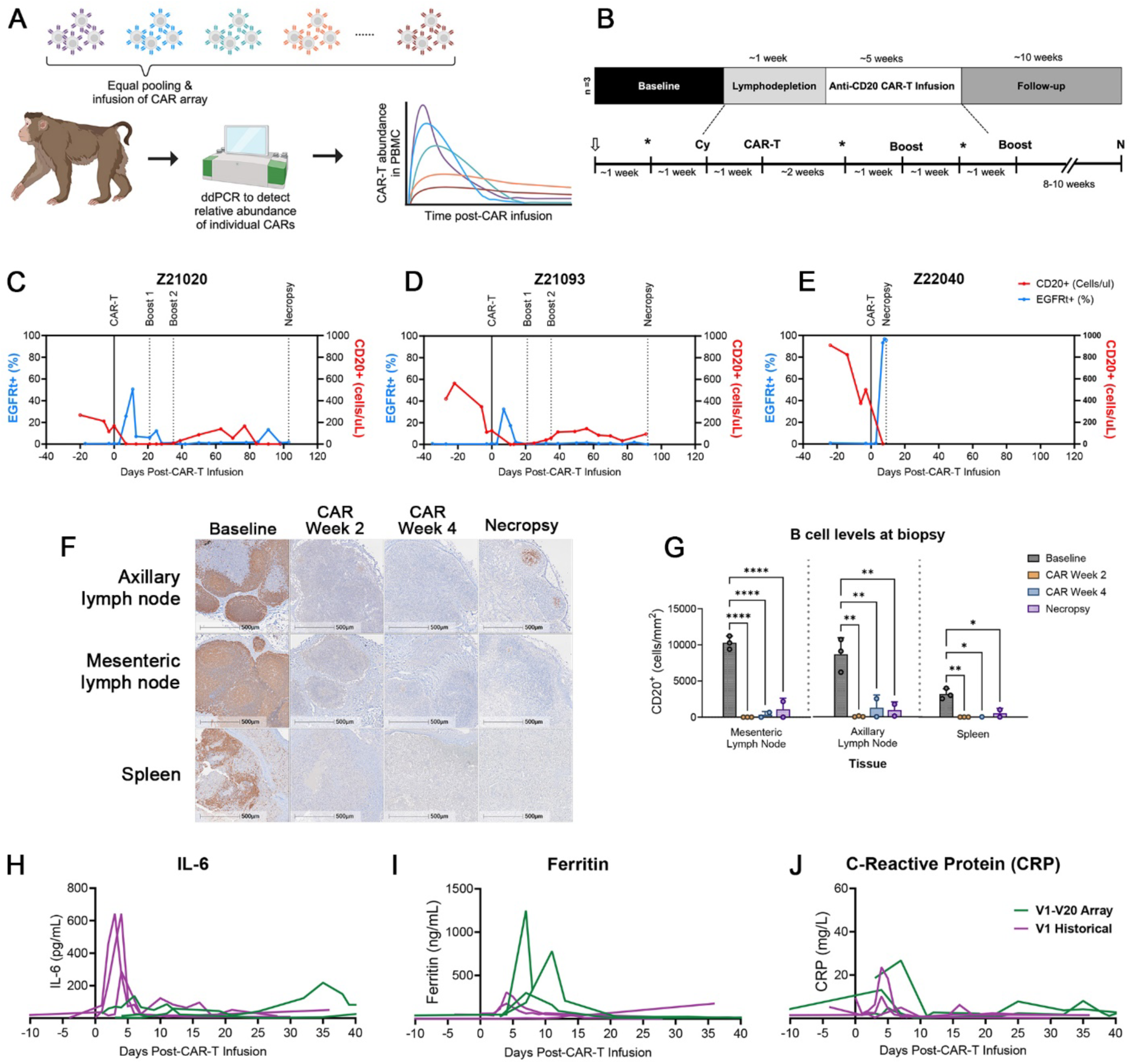
Infusion of the pooled CAR-T cell array drives robust expansion, durable B-cell depletion, and distinct inflammatory dynamics. (A) CAR-T variants were manufactured *ex vivo*, pooled in equal proportions, and infused into conditioned autologous NHPs. CAR variant abundance was tracked longitudinally using ddPCR. (B) Animals underwent leukapheresis (arrow) followed by cyclophosphamide (Cy) conditioning. Pooled CAR-T products were infused (CAR-T), and two K562-CD20 antigen boosts (Boost) were administered at weeks 3 and 5 post-infusion. Tissue samples (*) were collected at -2, 2, and 4 weeks relative to CAR-T cell infusion. Necropsy (N) occurred 9-102 days after CAR infusion. (C-E) Longitudinal analysis of CAR-T cell expansion (blue) and B-cell depletion (red) in PBMCs, for animal ID (C) Z21020, (D) Z21093, and (E) Z22040. (F) Representative immunohistochemistry (IHC) images from Z21020 showing CD20^+^ B cells (brown) in lymphoid tissues at indicated timepoints. (G) Quantification of CD20^+^ B cells from IHC across timepoints, with each data point representing an individual animal. (H-J) Longitudinal quantification of serum (H) IL-6, (I) ferritin, and (J) C-Reactive protein levels in the pooled CAR array cohort (V1-V20 CAR array) compared to a comparable historical single-CAR study that infused V1 (V1 Historical). Statistical significance for (G) was determined using one-way ANOVA with Tukey’s correction for multiple comparisons. \**P* < 0.05, \*\**P* < 0.005, \*\*\**P* < 0.0005, \*\*\*\**P* < 0.00005.

Total CAR-T cells expanded significantly after infusion, constituting 30-96% of peripheral CD3^+^ T cells, with peak levels observed between 7-11 days post-infusion (**Figure 5C-E**). CAR-T expansion corresponded with rapid B-cell depletion in peripheral blood (**Figure 5C-E**) and substantial B-cell follicle loss in lymph nodes and spleen, as confirmed by immunohistochemistry (**Figures 5F-G**). At day 25, we detected a modest secondary CAR-T expansion following a single infusion of K562-CD20 in Z21020 (**Figure 5C**). Z22040 exhibited extreme CAR-T expansion within the first week (92-96% of CD3^+^ T cells) and developed immune effector cell-associated neurotoxicity syndrome (ICANS) despite intensive supportive care (**Supplemental Table 1**), requiring an end of study at day 9. For Z21020 and Z21093, B-cells recovered between 30- and 40-days post-infusion, although levels remained low through study completion (**Figure 5G**). Z21020 and Z21093 developed gut infections approximately 80 days post-infusion that did not respond to antibiotic treatment. Based on clinical signs and low levels of CAR^+^ cells at these timepoints, these animals ended study at days 92 and 102.

Collectively, these data indicated increased CAR-T activity in NHPs receiving the pooled V1-V20 CAR array, relative to a previously-published cohort that received a single infusion of V1.^40^ Intriguingly, inflammatory biomarkers in serum were distinct between these cohorts. Unlike the historical V1 cohort, which exhibited a significant IL-6 spike after CAR-T infusion, the pooled CAR array cohort showed no IL-6 elevation (**Figure 5H**) but had markedly higher ferritin levels (**Figure 5I**). C-Reactive protein (CRP) levels were comparable (**Figure 5J**). Multiplex plasma cytokine analyses during peak CAR expansion confirmed reduced IL-6 levels, as well as decreased IFNγ and elevated IL-2 in the CAR array cohort (**Supplemental Figure 5A**, *P* < 0.05). We also performed lumbar punctures to collect CSF and observed distinct cytokine trends, with CAR array NHPs exhibiting increased IL-6, IFNγ, and IL-18 compared to the historical cohort (**Supplemental Figure 5B**). Collectively, the pooled CAR-T array demonstrated robust CAR-T expansion, durable B-cell depletion, and distinct, compartment-specific inflammatory dynamics, including muted IL-6 responses, elevated ferritin, and altered cytokine profiles.

### MyD88-CD40 CARs preferentially expand, traffic, and persist *in vivo*

We surmised that the increased CAR activity, toxicity, and divergent inflammatory profiles in CAR array animals relative to our historical V1 cohort were driven by distinct CAR variant(s) within the array. Consistent with this hypothesis, longitudinal variant-specific ddPCR revealed significant enrichment of MyD88-CD40 CAR-T cells (V13-V16) in PBMCs across all timepoints and NHPs (**Figure 6A-D**, *P* < 0.005). Likewise, MyD88-CD40 CAR-T cells were strikingly enriched in lymph nodes, spleen, bone marrow, BAL, and CSF samples collected within 2 weeks of CAR-T infusion (**Figure 6E**, *P* < 0.005). Among these, V14 (CD28 hinge/transmembrane domain) was most abundant during the primary expansion phase (**Figure 6F**) and clustered distinctly from other MyD88-CD40 variants via PCA (**Figure 6G**). 4-1BB CARs showed some early expansion and modest tissue trafficking, but did not persist beyond two weeks. Neither CD28 nor BAFF-R CARs showed appreciable expansion in peripheral blood or tissues. MyD88-CD40 CARs persisted in blood and tissues until necropsy, with marked enrichment in BAL and lung tissue samples (**Figure 7A**). At necropsy, V13, V14, and V15 showed significant enrichment across various tissues (**Figure 7B**), and PCA revealed these variants clustered distinctly from each other and the rest of the CAR array (**Figure 7C**). Our NHP study demonstrates that MyD88-CD40 domains drive superior CAR-T expansion, persistence, and tissue trafficking *in vivo*, significantly outperforming conventional 4-1BB and CD28 CAR-T cells.

**Figure 6.**
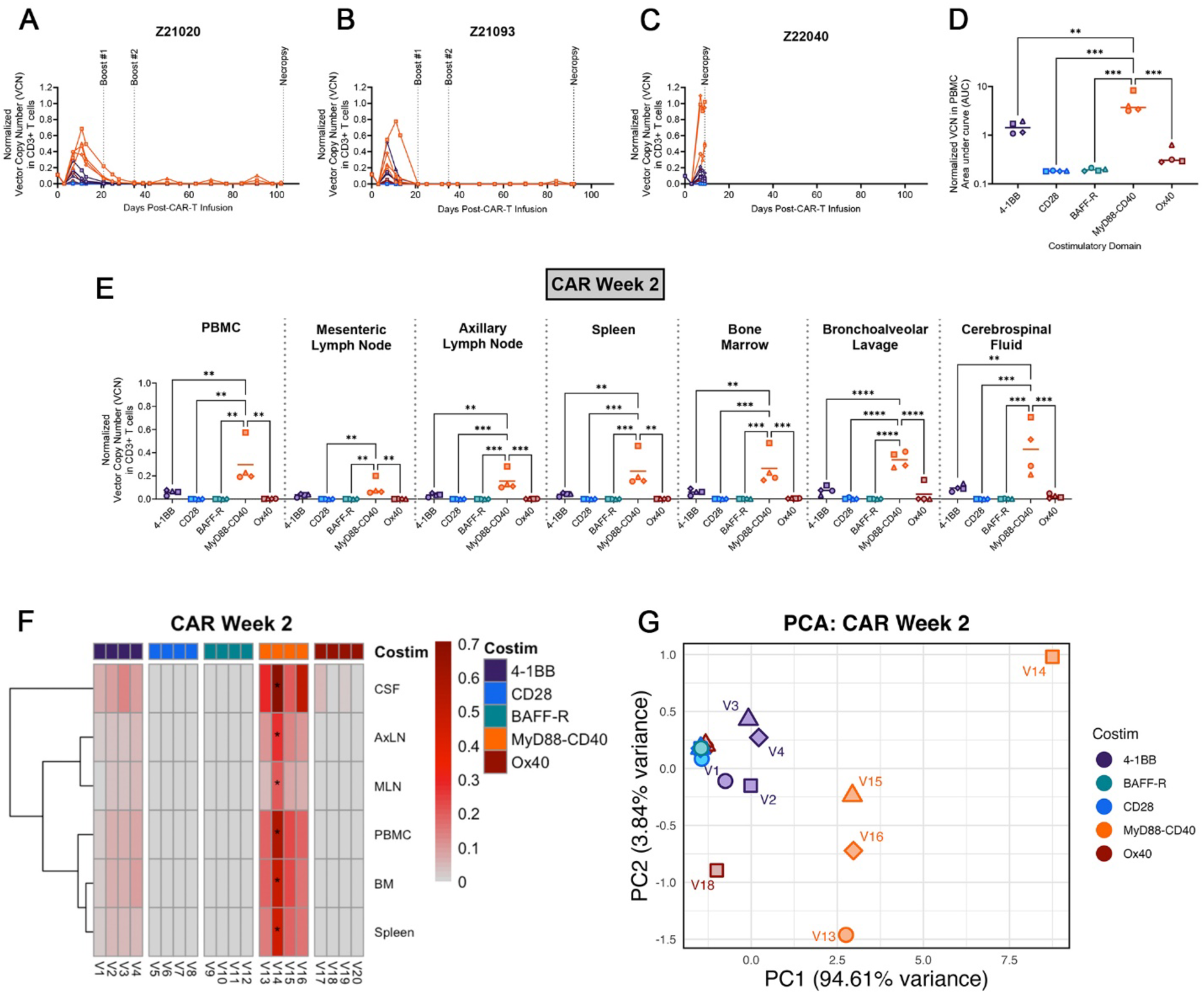
MyD88-CD40 CAR-T cells exhibit superior proliferation, durability, and tissue trafficking *in vivo*. Genomic DNA was extracted from blood and tissue samples to quantify the relative abundance of each CAR variant by ddPCR. Colors group each CAR by costimulatory domain; distinct hinge/transmembrane domains within each group are shown as different shapes as in Table 1. (A-C) Normalized vector copy number (VCN) of CAR variants from animal ID (A) Z21020, (B) Z21093, and (C) Z22040 PBMCs. (D) AUC of normalized VCN for CAR variants for Z21020 and Z21093. Each datapoint represents the average AUC from 0 to 92 days. (E) Normalized VCN from tissues collected between 9- and 14-days post-infusion, with each datapoint representing the average across three NHPs (*n* = 3 pigtail macaques), except for V17 (*n* = 2). (F) Normalized VCN from tissues collected between 9- and 14-days post-infusion, as in (E), displayed as a heatmap for statistical analysis by one-sample Z test with Benjamini-Hochberg correction to identify specific significantly enriched variants. Stars indicate significant variant abundance. (G) Data from (D-E) was compiled, Z-normalized and analyzed via PCA. Statistical significance for (D-E) was determined using one-way ANOVA with Tukey’s correction for multiple comparisons. \**P* < 0.05, \*\**P* < 0.005, \*\*\**P* < 0.0005, \*\*\*\**P* < 0.00005.

**Figure 7.**
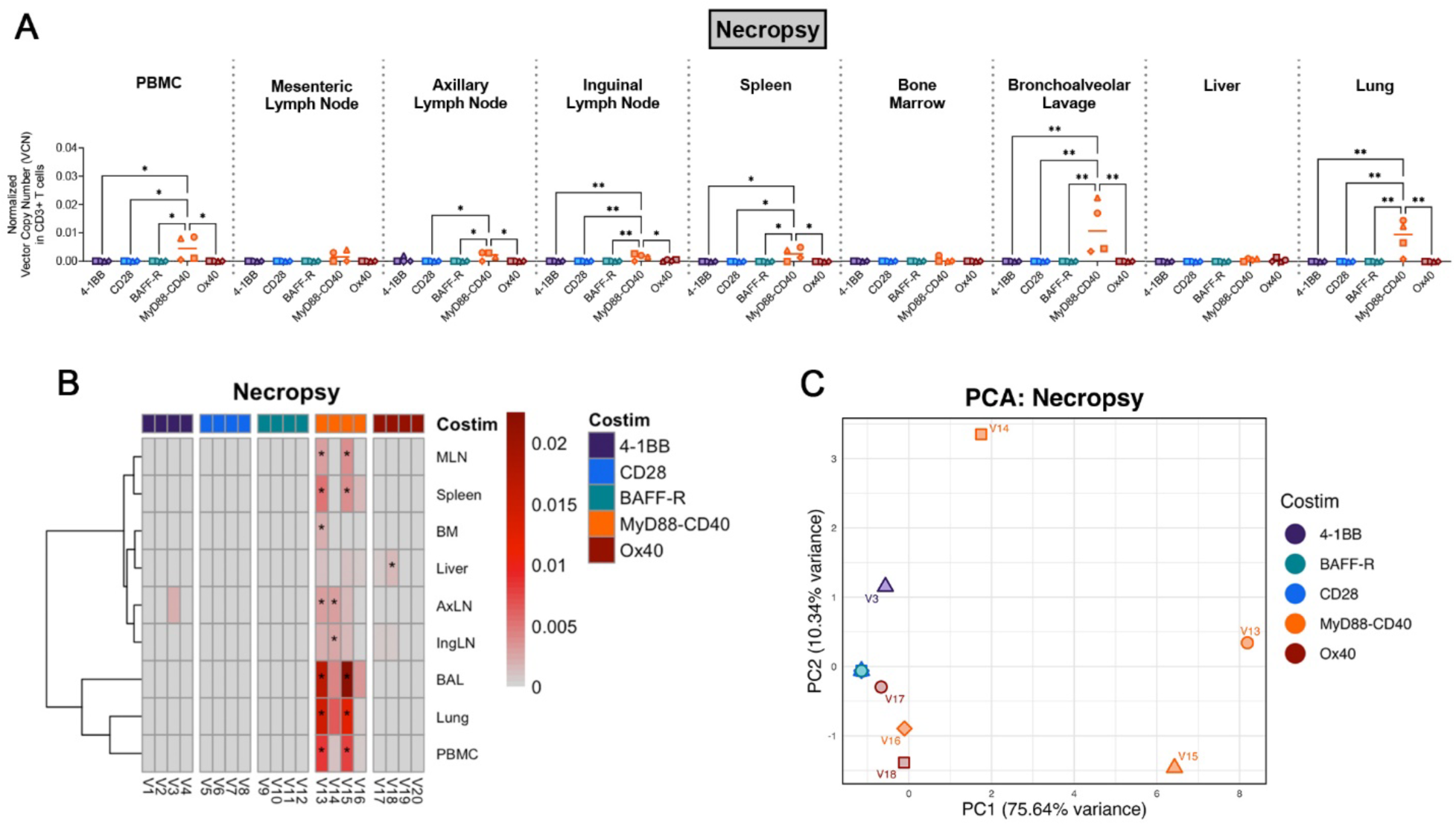
Necropsy analysis reveals durable persistence of MyD88-CD40 CAR-T cells. (A) Normalized VCN from tissues collected at necropsy. Each datapoint represents the average across two NHPs (Z21020 and Z21093), except for V17, where only one biological replicate (Z21093) was available. (B) Data from (A), displayed as a heatmap for statistical analysis by one-sample Z test with Benjamini-Hochberg correction to identify significantly enriched variants, indicated with stars. (C) Compiled data from (A) were Z-normalized and analyzed via PCA. Statistical significance in (A) was determined using one-way ANOVA with Tukey’s correction for multiple comparisons. \**P* < 0.05, \*\**P* < 0.005, \*\*\**P* < 0.0005, \*\*\*\**P* < 0.00005.

## DISCUSSION

While CAR-T cell therapy has redefined the treatment landscape for numerous B-cell malignancies, achieving long-term efficacy remains a challenge, in part due to suboptimal CAR-T cell persistence *in vivo*. Addressing this limitation will be paramount for unlocking the full potential of CAR-T therapies, not only in the context of B-cell malignancies but also for expanding their application to solid tumors, infectious disease, and autoimmunity.^40,57^ We developed a novel *in vivo* CAR competition assay in NHPs to simultaneously evaluate the behavior of multiple CAR designs in an immunocompetent background. We identified specific CAR architectures, notably the MyD88-CD40 costimulatory domain, that drive enhanced CAR-T cell proliferation, persistence, and tissue trafficking.

In primary cell-based assays, MyD88-CD40 CAR-T cells exhibited four key features: (1) enhanced activation, (2) tonic signaling, (3) distinct cytokine secretion, and (4) resistance to exhaustion following chronic antigen stimulation. These characteristics likely supported the robust *in vivo* proliferation of MyD88-CD40 CARs. MyD88-CD40 V14, which featured a CD28 hinge and transmembrane domain, demonstrated the strongest proliferation *in vivo*, consistent with prior reports that the CD28 transmembrane domain enhances CAR activity and antigen sensitivity.^33^ This feature may have supported the sustained persistence of MyD88-CD40 CAR-T cells even as B-cells declined. Importantly, other groups have similarly identified MyD88-CD40 as a superior costimulatory domain in pooled screening approaches, though predominantly in liquid tumor models, suggesting the efficacy of this domain may vary with tumor type and the associated microenvironment.^58^ This group also linked MyD88-CD40 to a hyperactivation and tonic signaling phenotype, reinforcing the reproducibility of our findings.

The impact of tonic signaling on CAR-T cell efficacy remains controversial. Many groups have linked tonic signaling to scFv stability and characterized it as a detrimental feature associated with excessive activation, terminal differentiation, and exhaustion.^59,60^ Others have noted that modest tonic signaling may improve CAR-T persistence and anti-tumor function.^61,62^ In our study, we maintained a consistent anti-CD20 scFv across all CAR designs, allowing us to attribute the tonic signaling primarily to intracellular domains. Notably, the tonic signaling we observed in MyD88-CD40 CARs did not lead to exhaustion; instead, these CARs displayed a robust resistance to exhaustion, a unique feature of this domain. This lack of exhaustion, combined with tonic signaling, likely synergized to enhance MyD88-CD40 CAR-T activity and durability *in vivo*.

The unique cytokine profile of MyD88-CD40 CAR-T cells offers key insights into their mechanisms of action and implications for next-generation CAR-T therapies. IL-6 is a hallmark of CAR-T expansion and CRS observed in patients and in prior NHP studies with 4-1BB CARs.^39,40,63^ The minimal IL-6 elevation we observed in MyD88-CD40 CAR-T cells both *ex vivo* and *in vivo* suggests that the characteristic IL-6 spike is primarily driven by conventional costimulatory domains like 4-1BB and CD28. As novel CAR architectures advance to the clinic, our findings underscore the importance of considering domain-specific toxicities. For example, our data indicate that IL-6 assays would be insufficient to diagnose CRS in patients receiving MyD88-CD40 CAR-T cells. MyD88-CD40 CAR-T cells also exhibited significantly reduced IL-10 secretion *ex vivo*. Given IL-10’s potent anti-inflammatory properties and critical role in regulating immunological homeostasis,^64^ its diminished secretion may have facilitated the robust expansion observed *in vivo*. These results align with emerging interest in combining CAR-T cells with IL-10 blockade, which has been shown to reinvigorate CAR-T activity in preclinical models.^65,66^ Rather than IL-6, MyD88-CD40 CAR-T cells secreted significantly increased IL-13 *ex vivo*, a cytokine associated with Th2-mediated immune responses and allergic inflammation.^62^ This inflammatory signaling may have supported the hyperactivation and tonic signaling observed in these CARs, suggesting that IL-13 secretion could be harnessed in armored CAR-T therapies to enhance activation and persistence.

Our pooled *ex vivo* stimulation assays, which were intended to simulate the pooled *in vivo* competition assay, revealed superior expansion of CD28 and 4-1BB CARs, and almost immediate depletion of MyD88-CD40 CARs. Importantly, these results were not replicated *in vivo*, where we saw striking expansion of MyD88-CD40 CAR-T cells and near-complete absence of CD28 CAR-T cells. This discrepancy may potentially be attributed to paracrine signaling factors present in the *ex vivo* setting that supported the expansion of these CAR designs. Our findings caution against overreliance on artificial *ex vivo* pooled screening approaches and emphasize the importance of evaluating CAR-T cell function in physiologically relevant preclinical models. We also validated the value of the NHP model in identifying acute toxicities, including those associated with MyD88-CD40 domains. In our study, one NHP experienced uncontrolled CAR-T expansion, suggesting that hyperactivation, tonic signaling, and absent regulatory mechanisms may also contribute to safety concerns. Similar toxicities have been observed clinically; in an early trial evaluating PSCA-specific CAR-T cells incorporating an inducible MyD88-CD40 switch for prostate cancer, dose-limiting toxicities led to trial discontinuation.^30^ While biological activity was observed, these findings are consistent with prior data indicating that MyD88-CD40 CARs may be more effective in liquid tumors than in solid tumors.^58^ Our results underscore the importance of immunocompetent NHP models in recapitulating key CAR-mediated toxicities observed in the clinic, which are rarely captured by murine models. Future NHP studies should focus not only on the continued development of novel CAR architectures but also on identifying prognostic markers of toxicity and devising strategies to mitigate the hyperactivity observed in MyD88-CD40-containing CARs, such as cis-acting safety switches.^67^

In summary, our study highlights the power of the NHP model as a platform for pooled screening of CAR designs, enabling the identification of superior architectures such as MyD88-CD40. The exceptional proliferation, persistence, and trafficking conferred by this domain may address fundamental challenges in CAR-T therapies, including limited persistence. The unique features associated with MyD88-CD40, including enhanced activation, tonic signaling, distinct cytokine secretion, and resistance to exhaustion, highlight key attributes that drive MyD88-CD40 potency and provide valuable insights that could be leveraged in the design of next-generation therapies.

## Acknowledgements

We thank Helen Crawford for assistance in preparing the manuscript. All NHP-related work was conducted at the Washington National Primate Research Center (WaNPRC), with additional support from Dr. Robert Murnane (veterinary pathology) and Solomon Wangari, Britni Curtis, Joel Ahrens, and Naoto Iwayama for tissue collection and laparoscopic procedures. We also thank the Fred Hutchinson Cancer Center Vector Production core, including Logan Hargis, Zach Burger, and Dr. Megha Gupta, for their work on lentiviral vector production; the WaNPRC Virology and Immunology core (WaNPRC V&IC), led by Dr. Sandra Dross, for cell subset analysis; and the University of Washington Laboratory Medicine Research Testing Service, including Dr. Chihiro Morishima and Jacquie Chee. We are grateful to Drs. Michael Jensen, James Riley, and Brian Till for providing critical reagents.

## Author contributions

L.H.M., H-P.K., K.R.J., and C.W.P. conceived and designed the study; L.H.M. manufactured the CAR-T cell products; L.H.M., E.J.C., and S.M.D. performed functional assays on CAR-T cell products; V.N., M.H., S.H., C.L., K.C., C.W., and E.W. provided care for NHPs; L.H.M., E.J.C., S.M.D., and T.E. collected and processed longitudinal samples and performed flow cytometry assays, which were analyzed by L.H.M.; L.H.M. extracted genomic DNA; H.Z. and L.S. performed ddPCR assays, which were analyzed by L.H.M.; C.E.S. performed and analyzed IHC; L.H.M. and C.W.P. wrote the manuscript, which was reviewed by all authors.

