## Supplemental Data for "Pooled CAR-T screening in nonhuman primates identifies designs with enhanced proliferation, trafficking, and persistence"

### LIST OF SUPPLEMENTAL ITEMS

1. **Supplemental Methods**
2. **Supplemental Table 1.** Overview of nonhuman primates infused with pooled CAR-T cell products.
3. **Supplemental Table 2.** List of antibodies used.
4. **Supplemental Figure 1.** Antigen-specific cytokine secretion and cytotoxicity profiles across CAR-T cell variants.
5. **Supplemental Figure 2.** Validation of ddPCR specificity and sensitivity.
6. **Supplemental Figure 3.** Pooled *ex vivo* chronic stimulation assay reveals donor-specific expansion of CAR-T cell variants
7. **Supplemental Figure 4.** Composition of pooled infusion products by variant.
8. **Supplemental Figure 5.** Cytokine profiles in plasma and cerebrospinal fluid (CSF) during acute CAR-T cell expansion.

### Supplemental Methods

#### Construction of CD20-targeted CAR array

The CAR array was designed to span all possible combinations of hinge (CD8, CD28), transmembrane (CD8, CD28) and costimulatory (4-1BB, CD28, BAFF-R, MyD88-CD40, Ox40) domains, with common anti-CD20 short chain variable fragment (scFv) and CD3ζ activation domains. The CAR array was codon-optimized for expression in *Macaca nemestrina* (pigtail macaques) and synthesized individually as gBlocks (Integrated DNA Technologies). We included a co-expressed marker (truncated epidermal growth factor receptor, EGFRt) downstream of the CAR, separated by a T2A sequence, as a surrogate marker for detecting CAR-transduced T cells. The CAR array was cloned into a self-inactivating lentivirus backbone (vector plasmids kindly provided by Dr. Michael Jensen) and sequence verified via next-generation sequencing (Plasmidsaurus).

#### ddPCR vector copy number (VCN) calculation

For all ddPCR reactions, 1–10 μL of gDNA was used as input, and raw ddPCR counts are reported as counts per μL of gDNA sampled. The total T cells in each sample were estimated by calculating the total cell genomes (based on MRPP30 counts, representing all cells) and subtracting TRD counts (representing non-T cells).[1] This difference was then divided by 2 to account for the two copies of each gene per genome:

$$\text{Total T cells}\ \left(\frac{\text{genomes}}{\mu\text{L}}\right) = \frac{\text{MRPP30}\ \left(\frac{\text{counts}}{\mu\text{L}}\right) - \text{TRD}\ \left(\frac{\text{counts}}{\mu\text{L}}\right)}{\left(\frac{\text{2 copies}}{\text{genome}}\right)}$$

The vector copy number (VCN) for individual CAR variants in both unmixed and pooled samples was calculated as the number of CAR variant counts per total T cell genomes in each sample:

$$\text{CAR variant VCN} \left(\frac{\text{copies}}{\text{T cell genome}}\right) = \frac{\text{CAR variant} \left(\frac{\text{counts}}{\mu\text{L}}\right)}{\text{Total T cells} \left(\frac{\text{genomes}}{\mu\text{L}}\right)}$$

**Normalization of ddPCR data from pooled assays**

In our *ex vivo* and *in vivo* experiments involving pooled CAR variants, we applied normalization to account for minor discrepancies in the initial composition of CAR variants in the pool. These discrepancies could arise from imprecisions in flow cytometry quantification of CAR marking prior to pooling, pipetting errors during pooling, or differences in vector copy number (VCN) integrations across CAR variant transductions. To address this, we normalized longitudinal ddPCR counts by the relative composition of CAR variants in the initial pooled sample. For example, if a particular CAR variant was found to be over-represented by a factor of 2 in the initial pool via ddPCR, we divided subsequent ddPCR counts for that variant by 2 to correct for this initial bias. The normalization process involved the following steps:

1. We calculated the relative abundance of each CAR variant ("CAR variant VCN abundance ratio") in the <u>initial pooled sample</u>:

$$\text{CAR variant VCN abundance ratio} = \frac{\text{CAR variant VCN} \left(\frac{\text{copies}}{\text{T cell genome}}\right)}{\begin{array}{c}\text{Average of all CAR}\\ \text{variant VCNs in pooled sample}\end{array} \left(\frac{\text{copies}}{\text{T cell genome}}\right)}$$

2. Next, raw CAR variant ddPCR counts from subsequent *ex vivo* or *in vivo* timepoints were normalized by the CAR variant VCN abundance ratio in the initial pooled sample:

$$\text{Normalized CAR variant} \left(\frac{\text{counts}}{\mu\text{L}}\right) = \frac{\text{CAR variant} \left(\frac{\text{counts}}{\mu\text{L}}\right)}{\begin{array}{c}\text{CAR variant VCN abundance ratio}\\ \text{in initial pooled sample}\end{array}}$$

3. We then calculated the normalized CAR variant VCN in total T cells for each CAR variant by dividing the normalized CAR variant counts to the measured total T cell counts from the same sample:

$$\text{Normalized CAR variant VCN}\left(\frac{\text{copies}}{\text{T cell genome}}\right) = \frac{\text{Normalized CAR variant}\left(\frac{\text{counts}}{\mu\text{L}}\right)}{\text{Total T cells}\left(\frac{\text{genomes}}{\mu\text{L}}\right)}$$

**Quantifying pooled CAR ddPCR sensitivity**

To evaluate ddPCR sensitivity, we simulated conditions of low CAR abundance. We manufactured a subset of 14 CAR variants using our standard CAR-T manufacturing, resulting in CAR-T cell variants ranging from 15–64% CAR expression, as measured by flow cytometry. Genomic DNA (gDNA) was extracted from each variant and pooled equally by mass (1:1 ratio). The pooled gDNA was then diluted with wildtype ($CAR^-$) gDNA (wt gDNA) at six levels: 0%, 10%, 50%, 90%, 96%, and 99% wt gDNA, creating a dilution series with varying CAR variant content. To confirm the ability of ddPCR to reliably detect each of the 14 variants across the dilution series, we applied variant-specific ddPCR assays to quantify variant abundance. To account for differences in transduction efficiency and vector copy number (VCN) among the variants, we also measured individual CAR variant VCNs from gDNA isolated from unmixed CAR-T variants. We plotted observed ddPCR counts against expected counts, where the expected ddPCR counts were calculated as follows:

1. The vector copy number (VCN) for <u>unmixed CAR variants</u> was calculated by dividing the CAR variant counts by the MRPP30 counts, then adjusting for the two copies of MRPP30 per genome:

$$\text{CAR variant VCN}\left(\frac{\text{copies}}{\text{genome}}\right) = \left(\frac{\text{CAR variant}\left(\frac{\text{counts}}{\mu\text{L}}\right)}{\text{MRPP30}\left(\frac{\text{counts}}{\mu\text{L}}\right)}\right)\cdot\left(2\ \text{MRPP30}\left(\frac{\text{copies}}{\text{genome}}\right)\right)$$

2. We quantified the number of genomes per μL in the six dilution samples by measuring MRPP30 counts using ddPCR and dividing by 2 to account for the two copies of MRPP30 per genome:

$$\left(\frac{\text{Genomes}}{\mu\text{L}}\right)\text{ in dilution series} = \frac{\text{MRPP30}\left(\frac{\text{counts}}{\mu\text{L}}\right)}{2\ \text{MRPP30}\left(\frac{\text{copies}}{\text{genome}}\right)}$$

3. We then calculated the expected CAR variant copies per μL by multiplying the CAR variant VCN by $\left(\frac{1}{14}\right)$, to account for the 14 CAR variants pooled, and by the number of genomes per μL:

$$\text{Expected CAR variant}\left(\frac{\text{copies}}{\mu\text{L}}\right) = \text{CAR variant VCN}\left(\frac{\text{copies}}{\text{genome}}\right)\cdot\left(\frac{1}{14}\right)\cdot\left(\frac{\text{genomes}}{\mu\text{L}}\right)$$

4. Next, we adjusted the expected CAR variant copies per μL to account for the applied dilution factor by multiplying the expected copies per μL by the fraction of wt gDNA vs $CAR^+$ gDNA in the sample:

$$\text{Adjusted CAR variant}\left(\frac{\text{copies}}{\mu\text{L}}\right) = \text{CAR variant}\left(\frac{\text{copies}}{\mu\text{L}}\right)\cdot(\text{1-fraction wt gDNA})$$

5. To match the observed sample measurements, we multiplied the expected variant copies per μL by 10 to account for the 10 μL of gDNA assayed. The expected variant copies per 10 μL were then plotted against the observed ddPCR counts per 10 μL of gDNA from the dilution series.

**CRS management and serum cytokine assays**

Animals received tocilizumab (8 mg/kg per dose), anakinra (1 mg/kg per dose), and dexamethasone (0.5 - 2 mg/kg per dose) under clinician direction to prevent and treat CRS (**Supplemental Table 1**). Z22040 experienced ongoing symptoms of immune effector cell-associated neurotoxicity syndrome (ICANS), and was given Keppra (15 - 60 mg/kg), furosemide (2 mg/kg), and diazepam (0.5 mg/kg) to prevent and treat seizure activity. Blood samples were analyzed by the University of Washington Clinical Laboratory for C-reactive protein (CRP), ferritin and IL-6.

**Immunohistochemistry**

Immunohistochemistry was performed as previously described.[2] Briefly, 5-μm tissue sections mounted on glass slides were deparaffinized in xylene and rehydrated through a series of graded ethanol to distilled water solutions. Heat-induced epitope retrieval (HIER) was performed with the antigen retrieval citrate buffer (pH 6.0) in a NxGen decloaking chamber (Biocare Medical) at 110°C for 15 min, cooled for 20 min, and then rinsed twice in dd$H_2O$ and 1x TBS with 0.05% Tween 20 (TBS-T). Slides were incubated with blocking buffer (TBS-T with 0.25% casein) for 30 min at room temperature, rinsed with TBS-T, and incubated at room temperature for 1 hour with goat anti-CD20 (Invitrogen; cat. no. PA1-9024; at 1:800), which was diluted in blocking buffer. Endogenous peroxidases were blocked with 1.5% $H_2O_2$ in TBS-T for 5 min. Slides were then incubated with goat Polink-1 HRP (Origene). Slides were developed using Impact DAB (3,3′-diaminobenzidine; Vector Laboratories), washed in dd$H_2O$, counterstained with hematoxylin (Biocare Medical), mounted in Permount (Fisher Scientific), and scanned at 20x magnification. Quantitative image analysis was performed using HALO software (v.3.6.4134; Indica Labs) on lymph node or splenic sections from each animal. For CD20 quantification, the Multiplex IHC module v3.4.9 was used to detect $CD20^+$ cells, and quantification is presented as a proportion of total tissue ($CD20^+$ cells/$mm^2$). Manual curation was performed to confirm accurate annotations.

**Supplemental Table 1: Overview of nonhuman primates infused with pooled CAR-T cell products.**

| Animal ID (sex) | Weight* (kg) | Dosage ($CAR^+$ cells/kg) | Peak CAR expansion** | Days to B cell recovery† | Clinical observations during peak expansion | Supportive medications during peak expansion | Duration of study (days) |
|---|---|---|---|---|---|---|---|
| Z21020 (M) | 4.7 | $9 \times 10^6$ | 50.6% | 39 days | Fever, inappetence | - | 102 |
| Z21093 (M) | 4.1 | $9 \times 10^6$ | 32.5% | 32 days | Fever, inappetence, diarrhea, dehydration, itching | Benadryl, tocilizumab, anakinra, dexamethasone | 92 |
| Z22040 (M) | 3.7 | $9 \times 10^6$ | 96.8% | - | Fever, inappetence, tremors, altered consciousness, seizure activity | Benadryl, tocilizumab, anakinra, dexamethasone, diazepam, furosemide, Keppra | 9 |

* Weight on day of infusion

** Peak CAR expansion as a percentage of total $CD45^+$ $CD3^+$ T cells in PBMCs by flow cytometry

† Days to B cell recovery defined as the number of days until $CD20^+$ cell counts exceed 20 cells per μL.

**Supplemental Table 2: List of antibodies used**

| Target | Clone | Supplier | Assay |
|---|---|---|---|
| CD3 | SP34-2 | BD | Flow cytometry |
| CD45 | D058-1283 | BD | Flow cytometry |
| CD4 | L200 | BioLegend | Flow cytometry |
| CD8 | SK1 | BD | Flow cytometry |
| PD-1 | Z12.2H7 | eBioscience | Flow cytometry |
| TIGIT | MBSA43 | eBioscience | Flow cytometry |
| CD25 | BC96 | BioLegend | Flow cytometry |
| HLA-DR | G46-6 | BD | Flow cytometry |
| Ki67 | DX2 | BD | Flow cytometry |
| EGFRt | Hu1 | Biotechne | Flow cytometry |
| CD22 | S-HCL-1 | BD | Flow cytometry |
| CD28 | CD28.2 | BD | Flow cytometry |
| CD95 | DX2 | Invitrogen | Flow cytometry |
| CD20 | Polyclonal<br>(cat. no. PA1-9024) | Invitrogen | IHC |

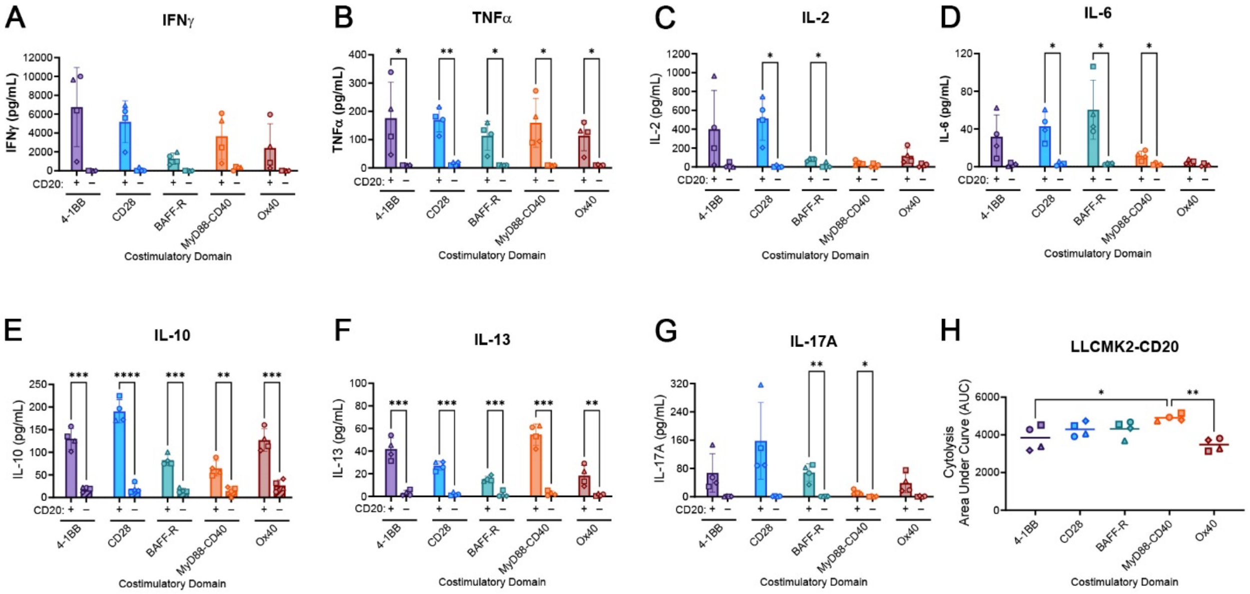


**Supplemental Figure 1. Antigen-specific cytokine secretion and cytotoxicity profiles across CAR-T cell variants.** (A-G) CAR-T variants were co-cultured with irradiated K562-CD20 or K562-wt cells. Supernatants were collected after 24 hours and analyzed for (A) IFNγ, (B) TNFα, (C) IL-2, (D) IL-6, (E) IL-10, (F) IL-13, or (G) IL-17A cytokine secretion. Antigen status of the target cells ($CD20^+$ or $CD20^-$) is indicated on the x-axis. (H) Area-under-the-curve (AUC) quantification of CAR variants' cytolysis of co-cultured LLCMK2-CD20 target cells, as outlined by methods depicted in Figure 3H-I. To compare cytokine secretion between $CD20^+$ and $CD20^-$ conditions, data in (D-F) were reanalyzed from Figure 3E-G using unpaired t-tests with Holm-Šídák correction. To assess antigen-specific cytotoxicity across costimulatory groups, LLCMK2-CD20 cytolysis data in (H) were reanalyzed from Figure 3I using one-way ANOVA with Tukey's correction. Each data point in (A-G) represents the mean of three biological replicates (*n* = 3 pigtail macaques), except for V17 (*n* = 2). (H) includes two replicates. Statistical significance: **P* < 0.05, ***P* < 0.005, ****P* < 0.0005, *****P* < 0.00005.

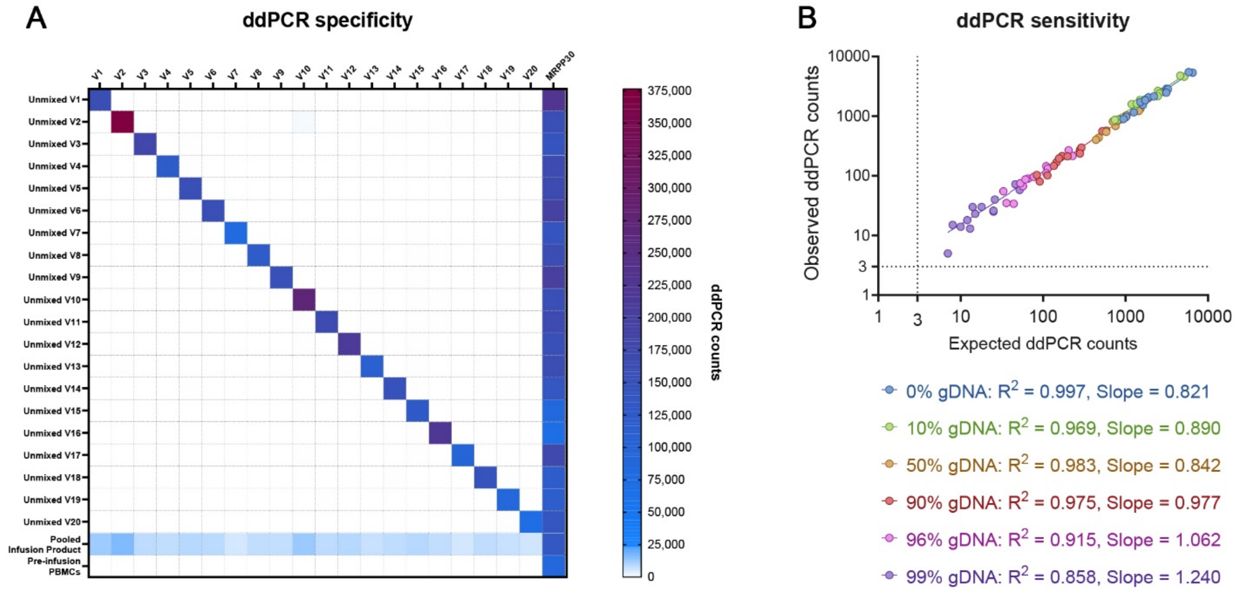


**Supplemental Figure 2. Validation of ddPCR specificity and sensitivity.** (A) Variant-specific ddPCR was used to analyze samples indicated on the y-axis, including genomic DNA (gDNA) from individually manufactured CAR-T variants (unmixed V1– unmixed V20), pooled CAR-T cell infusion products, and pre-infusion ($CAR^-$) PBMCs. Heatmap displays the average ddPCR counts across three biological replicates (*n* = 3 pigtail macaques), except for V17 (*n* = 2); white boxes indicate zero cross-variant background for each CAR-specific primer probe set (x-axis). The far-right column in the matrix shows a genomic control/housekeeping locus (macaque gene RPP30, MRPP30), used to quantify total genomes for each y-axis sample. (B) To evaluate ddPCR sensitivity, we simulated conditions of low CAR abundance. gDNA from 14 CAR-T variants was mixed in equal proportions by mass and serially diluted with wild-type gDNA (0%, 10%, 50%, 90%, 96%, 99%). Expected ddPCR counts, computed based on the measured VCN of the individual CAR-T variants and the applied dilution factors (see Supplemental Methods), were compared with observed counts. The assay reliably resolved expected 1:1 input ratios across all dilution levels, maintaining strong correlations between expected and observed counts ($0.858 < R^2 < 0.997$; $0.821 < \text{slope} < 1.240$). Individual CAR variants were consistently detected up to 99% wild-type gDNA spike-in levels, above the assay's positivity threshold of ≥3 ddPCR counts.

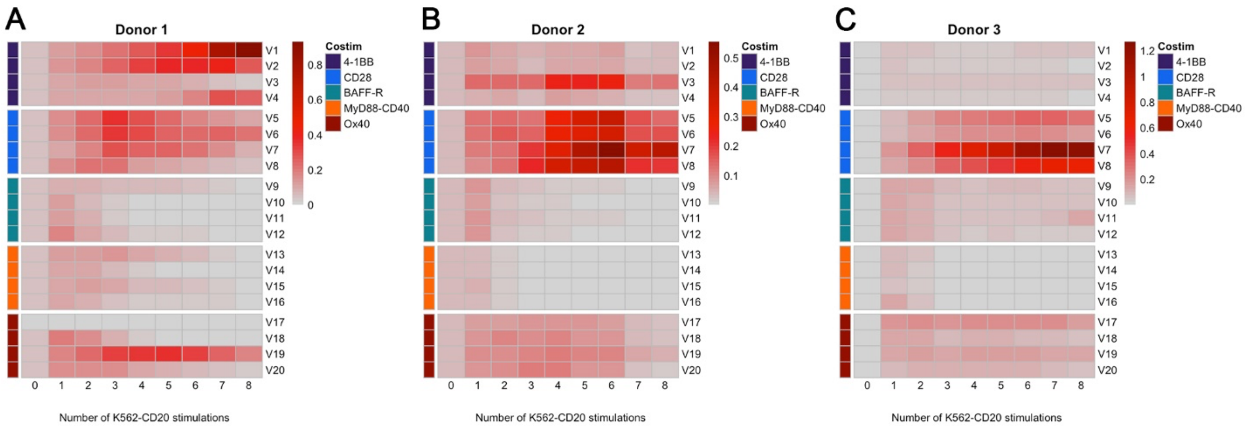


**Supplemental Figure 3. Pooled *ex vivo* chronic stimulation assay reveals donor-specific expansion of CAR-T cell variants.** (A-C) Equal numbers of CAR$^+$ cells from all CAR-T variants were pooled, subjected to chronic antigen stimulation, and analyzed for relative variant abundance using ddPCR. Heatmaps depict normalized vector copy number (VCN)—calculated by normalizing the measured VCN at each timepoint to the CAR Variant VCN abundance ratio in the initial pooled sample (see Supplemental Methods)—across three biological donors, as indicated above. Donors correspond to 3 pigtail macaques that were subsequently studied in *ex vivo* experiments. Donor 1: ID Z21020; Donor 2: ID Z21093; Donor 3: Z22040.

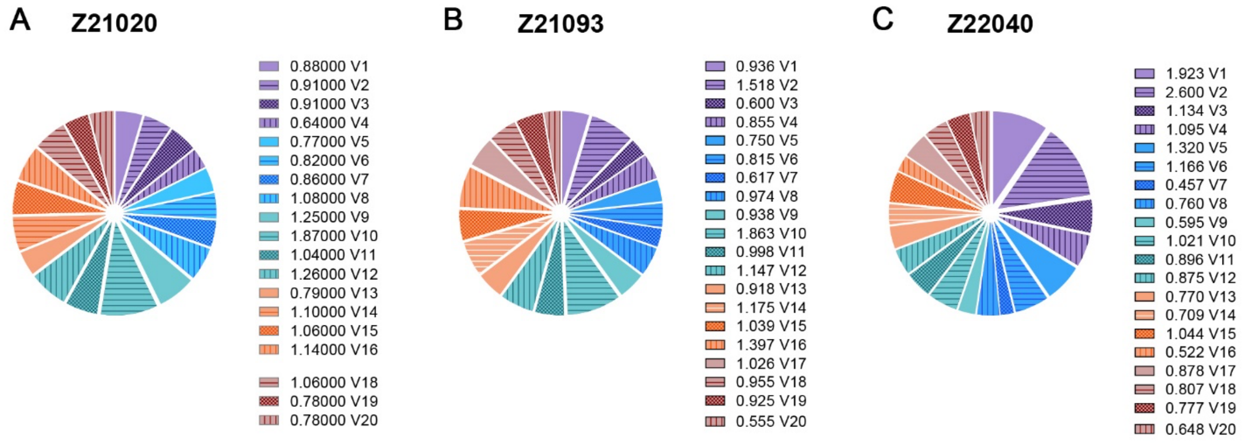


**Supplemental Figure 4**. **Composition of pooled infusion products by variant.** The relative abundance of each CAR variant in the pooled infusion product was quantified by ddPCR-based vector copy number (VCN) assay (see Supplemental Methods). Shown are data from animal IDs (A) Z12020, (B) Z21093, and (C) Z22040. Pie charts indicate each CAR variant with distinct colors/patterns. To assess the representation of each variant within the initial pooled sample, the VCN of each variant was normalized to the average VCN across all variants. The resulting CAR Variant VCN abundance ratio is shown for each CAR variant to the right of the pie chart. If all variants are precisely pooled, each would contribute equally with a relative ratio of 1. Z21020 did not receive V17 due to technical difficulties in manufacturing.

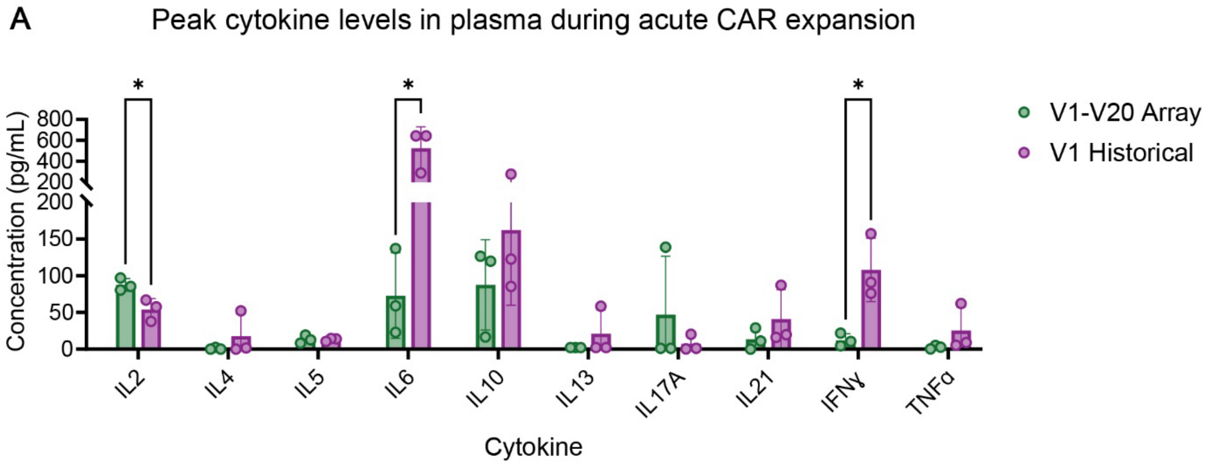


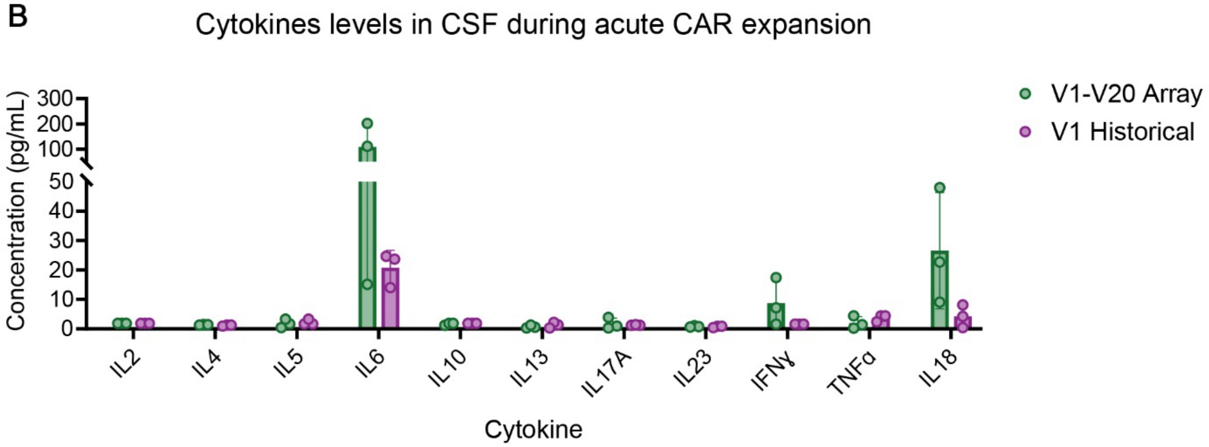


**Supplemental Figure 5. Cytokine profiles in plasma and cerebrospinal fluid (CSF) during acute CAR-T cell expansion.** (A) Peak cytokine levels in plasma during the acute phase of CAR-T cell expansion (days 3–13 post-infusion). (B) Cytokine levels in CSF samples obtained 9-14 days after CAR-T infusion. Cytokine concentrations for the pooled V1-V20 CAR array (green) were compared to a comparable historical single-CAR cohort that received V1 (purple). Each data point represents one animal (n = 3 pigtail macaques per cohort). All samples were assayed concurrently under identical conditions. Statistical significance was calculated using unpaired t-tests (*$P < 0.05$).
